# Bayesian Correlation is a robust similarity measure for single cell RNA-seq data

**DOI:** 10.1101/714824

**Authors:** Daniel Sanchez-Taltavull, Theodore J. Perkins, Noelle Dommann, Nicolas Melin, Adrian Keogh, Daniel Candinas, Deborah Stroka, Guido Beldi

## Abstract

**Assessing similarity** is highly important for bioinformatics algorithms to determine correlations between biological information. A common problem is that similarity can appear by chance, particularly for low expressed entities. This is especially relevant in single cell RNA-seq (scRNA-seq) data because read counts are much lower compared to bulk RNA-seq.

Recently, a **Bayesian correlation** scheme, that assigns low similarity to genes that have low confidence expression estimates, has been proposed to assess similarity for bulk RNA-seq. Our goal is to extend the properties of the Bayesian correlation in scRNA-seq data by considering 3 ways to compute similarity. First, we compute the similarity of pairs of genes over all cells. Second, we identify specific cell populations and compute the correlation in those populations. Third, we compute the similarity of pairs of genes over all clusters, by considering the total mRNA expression.

We demonstrate that Bayesian correlations are more reproducible than Pearson correlations. Compared to Pearson correlations, Bayesian correlations have a smaller dependence on the number of input cells. We show that the Bayesian correlation algorithm assigns high similarity values to genes with a biological relevance in a specific population.

We conclude that Bayesian correlation is a robust similarity measure in scRNA-seq data.

## 1 Introduction

Single cell RNA-seq (scRNA-seq) is one of the most recent advances in single cell technologies and it has been widely used to study multiple biological processes [64, 7, 21, 20, 12, 55, 23, 73, 51]. Standard bulk RNA-sequencing retrieves the average of RNA expression from all cells in a specific sample, thus providing an overall picture of the transcriptional activity at a given time point from a mixed population of cells. However, within the study of heterogeneous populations it is not possible to understand the contribution of individual cell types which is needed to dissect precise mechanisms. Single-cell RNA-seq (scRNA-seq) overcomes the limitations of bulk RNA-seq by sequencing mRNA in each cell individually, making it possible to study cells at a genome wide transcriptional level within heterogeneous samples. However, due to the small amount of mRNA sequenced within a cell, typically 80-85% of all genes remain undetected, a phenomenon known as dropout. This results in an incomplete picture of the mRNA expression pattern within a cell.

ScRNA-seq observations are biased because false zero counts occur in most genes because of the low amount of mRNA sequenced in each cell. There are several methods to correct for the dropout by imputing gene expression based on the gene expression of other cells. For example, MAGIC corrects gene expression with the gene expression of other cells modelled as a diffusion map [71]. scImpute corrects the dropout using similar cells and genes not affected by dropout [34]. SAVER uses a negative binomial to model gene expression in each cell and corrects dropout using the expression of other genes as predictors in a LASSO regression [24, 69]. All these methods follow the same principle by sacrificing part of the single cell structure of the data in order to obtain a better resolution of the different populations.

A similarity measure in mathematics is a function (with real values) that quantifies how similar two objects are. In the context of this project, we aim to determine similarity of genes in two distinct conditions. Several techniques use different notions of similarity to visualize data such as PCA or t-SNE. Several techniques exist to cluster data in scRNA-seq, such as Seurat [10], SCENIC [3] or Cell Ranger [78]. There are methods that use the notion of similarity to infer the gene regulatory dynamics. Some examples are SCENIC [3] or NetworkInference [13]. These techniques rely on data transformations and corrections of the dropout, but do not incorporate a notion of uncertainties in the measurements.

Assessing similarity between genes have previously been used in biology for biomarker discovery in cancer [39, 14], find patterns in gene expression [29] or to build gene expression networks [11, 68]. However, assessing similarity can be challenging since measurements of small populations with large uncertainties may lead to false correlations. If a gene’s expression is so low that it only registers zero or a few reads per cell, then its expression pattern across cells cannot be meaningfully related to that of other genes, there is simply too much uncertainty about the real expression levels of that gene. In a typical scRNA-seq dataset, the majority of genes may be in this situation, so that gene-gene correlation analysis is swamped with meaningless or spurious correlations.

Noise in gene expression measurements has been modeled and studied to identify differentially expressed genes [36, 31, 70]. Recently, uncertainties have been incorporated in methods to study differential expression in RNA-seq experiments [52] and a Bayesian scheme has been proposed to identify differentially expressed genes in scRNA-seq [63]. Noise is especially important in scRNA-seq because the low number of read counts. Therefore, methods to assess similarity in bulk RNA-seq may not be appropriate for scRNA-seq. Thus, methods need to be modified properly in order to maintain reproducibility. A simple solution is the removal of cells with a low number of read counts and low expressed genes, which is the currently used method of single-cell analysis [37]. However, there is not a systematic method to select a threshold and it highly depends on the population being studied.

In order to address limitations dependent on the noise, Bayesian statistics have been used to study biological processes [75, 74]. Recently, we have proposed a Bayesian correlation scheme to asses similarity between two entities in high-throughput sequencing (HTS) experiments [60, 56]. Such Bayesian correlation consider uncertainties in the measurements. And therefore, assign low values to correlations coming from low expressed genes using a prior belief and compute posterior belief based on data observation. Using this method, we have identified that a correct choice in the prior is crucial. Thus, we set out to develop new Bayesian methods to create better clustering algorithms than the ones currently available.

An additional consideration that needs to be addressed in scRNA-seq experiments is the number of sequenced cells. Simulation methods for scRNA-seq data is not a mature field and could not reproduce all the biological mechanisms present in an experiment. To study the effect of the number of cells on the reproducibility of the results, we sequenced parenchymal and non-parenchymal cells from a mouse liver. To test the sensitivity of the methods, 4 samples with an increasing number of input cells were explored (1000, 2000, 5000 and 10000 cells). To avoid biological noise, samples sequenced were from the same animal.

In this manuscript we show that Bayesian correlation is a robust similarity measure for pairs genes in single cell RNA-seq. We show that the reproducibility of Bayesian correlation is higher than the reproducibility of Pearson correlation. We show that the results obtained with Bayesian correlation have a small dependence to the number of cells, making the method suitable to study rare populations. Finally, we show that biologically relevant genes tend to appear more often in the top correlated pairs using Bayesian correlations.

## 2 MATERIALS AND METHODS

### 2.1 Mathematical Formulation of the Bayesian Correlation Method

After counting the aligned reads to exons and debarcoding the reads from cells, the output of a scRNA-seq experiment is the *n* × *m* unique molecular identifiers (UMI) matrix, *R*, where *n* is the number of genes and *m* the number of cells. Ideally we would correlate the true fraction of UMIs, *p_ie_*, of gene *i* in cell *e*. A trivial approximation is to normalize the data dividing every UMI by the total number of UMIs in that cell, that is 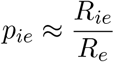. Bayesian schemes try to approximate *p_ie_* using a prior belief and compute posterior belief based on data observation.

Assume we have *R_ie_* UMIs of gene *i* in cell *e* and the total UMIs in that cell is *R_e_*. The empirical estimate is 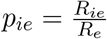. Then, Pearson correlation can be computed as

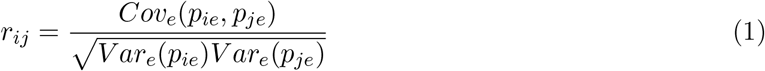

Whilst the Bayesian scheme would give us

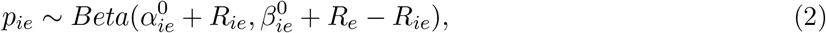

where 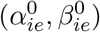 is the prior. In that scenario the covariance and variance are computed as

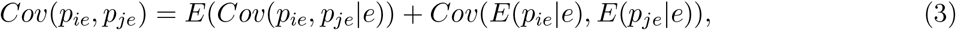

and

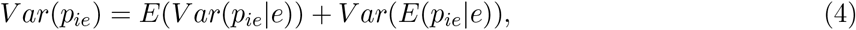

with

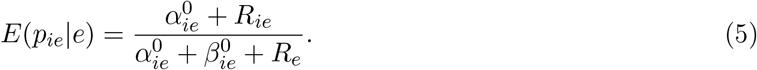

Then, the Bayesian Correlation is defined as

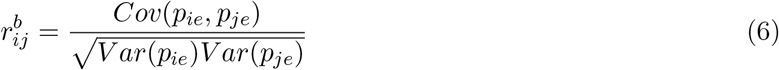

### 2.2 Cell isolation

Hepatocytes were isolated by a two-step collagenase perfusion. Animals were anesthetized (fentanyl 50ug/kg, midazolam 5mg/kg, medetomidine 500ug/kg, i.p.), immobilized in a supine position and the liver and portal vein exposed. The portal vein was cannulated with a 22G catheter and perfusion at 4ml/min with the buffers allowed to run to waste through an incision in the inferior vena cava. The liver was perfused with 10 ml of HBSS (Mg2+, Ca2+ free,10mM HEPES, pH7) followed by 25ml of HBSS containing EDTA (10mM HEPES pH7, 5mM EDTA). The EDTA was removed from the liver by perfusion with 10 ml of HBSS followed by digestion with 25 ml of HBSS containing collagenase (10mM HEPES, 1mM CaCl2, 0.5mg/ml collagenase IV, 0.01mg/ml collagenase 1A (Sigma)). The liver was then removed and the cells released by cutting the capsule and gentle agitation of the digested liver in stop buffer (HBSS, 10mM HEPES pH7, 5mM EDTA, 10mM citrate, 1% FBS) and passed through a 70*μ*m filter. The cell suspension was spun at 30G min to pellet most of the hepatocyte fraction, the supernatant collected and remnant cells pelleted at 250xG. This cell pellet was washed once in stop buffer then re-suspended in 20% isotonic Percoll and overlaid on a layer of 80% isotonic Percoll and spun at 500G for 10 minutes. The cells at the interface of the two Percoll layers were collected and washed in PBS, re-suspended in PBS and counted using a cell counter (BioRad TC 20).

### 2.3 Library preparation

scRNA-seq libraries were prepared from 1000, 2000, 5000 and 10000 cells using the Chromium Single Cell 3’ Library & Gel Bead Kit v3 (10xGenomics). Libraries were prepared according to the manufacturer’s protocol.

### 2.4 Sequencing

Sequencing was performed on a NovaSeq 6000 S2 flow cell. Read 1 consisted of 26 cycles (10XGenomics barcode plus UMI) followed by a single illumina i7 index read of 8 cycles and read 2 of 91 cycles to determine transcript-specific sequence information.

### 2.5 Read alignment

The function *cellranger count* from Cell Ranger was used to transform the fastq files with the parameter *expect-cells* set to 1000, 2000, 5000 or 10000. The reference genome was the mm10 available at Illumina Cell Ranger webpage. Next, we used *cellranger mat2csv* to generate the UMI matrix.

### 2.6 Data pre-processing

First, we created an SCE object with the function *SingleCellExperiment* from the R-package SingleCellExperiment.

The UMI matrix was filtered as follows: first, genes with 0 reads where excluded; second, cells with more than 15% of UMIs in mitochondrial genes were removed (mitochondrial gene list is included in suplementary materials). Cells with more than 25% UMIs in globin genes were removed. Finally, genes expressed in less than 2 cells were removed. Additionally, for the 5000-sample, a cell containing 110270 UMIs was considered an outlier and it was removed because the second cell with most UMIs had 26038 UMIs.

### 2.7 Dimensionality reduction and clustering

In order to cluster the data and find the different cell populations and their markers, we followed the procedure of Seurat [10]. The filtered UMI matrix was transformed into a Seurat Object with *CreateSeuratObject* with parameters *min.cells = 1* and *min.genes = 2*. We normalised the data with the R-function NormalizeData from Seurat with parameters: *normalization.method* = *“LogNormalize”* and *scale.factor = 10000*. Then, the data was scaled with the Seurat function *ScaleData* with parameter *vars.to.regress = c(“nUMI”)* The different clusters were identified using the Seurat function FindClusters with parameters *reduction.type = “pca”, dims.use = 1:8, resolution = 1.0, print.output = 0, save.SNN = TRUE*. t-SNE dimensionality reduction was done with the Seurat function RunTSNE with parameters: *dims.use = 1:8, do.fast = TRUE*. The different markers of each cluster were identified with the function *FindAllMarkers* with parameters *only.pos = TRUE, min.pct = 0.25, thresh.use = 0.25*

### 2.8 Drop-out correction

We corrected the dropout from the UMI matrix with the magic function form the R-package Rmagic with the parameters *genes* set equal to *“all_genes”* and default parameters. Any resulting negative expression was replaced with 0.

### 2.9 Notions of similarity

In this manuscript, we consider 3 notions of similarity between genes: **1-All cells correlation**: The correlation of gene *i* and gene *j* is the correlation coefficient using all cells. **2-Cluster correlation**: The correlation of gene *i* and gene *j* is the correlation coefficient using each cluster as condition, where the gene expression is the sum of the gene expression in all cells. That is, let *K* be the number of clusters, let 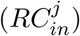 be the UMIs of gene *i* of cell *n* in cluster *j*. We transformed our *K* matrices in the reduced *n* × *k* dimensional matrix (*B_ij_*) as follows

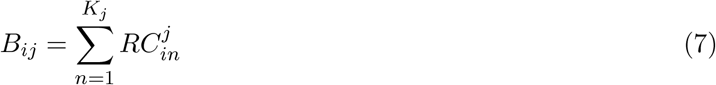

where *K_j_* in the number of cells in cluster *K*. The Bayesian correlation algorithm is applied to the bulk-like RNA-seq matrix (*B_ij_*). **3-In cluster correlation**: The correlation of gene i and gene j is the correlation coefficient using all cells in a specific cluster.

## 3 RESULTS

### 3.1 Bayesian correlation and Pearson correlation agreement increase with the number of cells

We studied the first notion of similarity: All cells correlation. To study the effect of the number of cells on the reproducibility of the results, the Bayesian correlation was computed and was compared with the Pearson correlation.

After the data processing, we ended up with four samples of 705, 1213, 2939 and 5520 cells. We refer to these samples as 1000-sample, 2000-sample, 5000-sample and 10k-sample.

Our first step was to compare our Bayesian similarity measure, using as a prior 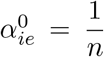 and 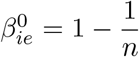 with Pearson correlation. In doing so, we split the samples into two, randomly assigning half of the cells to group *A* and the other half to group *B*. All pairwise correlations were independently computed for each group. In Fig. 1(a) we observed that the agreement between the two samples is higher using the Bayesian method. As the number of input cells increases, the agreement between the two groups increases. In Fig. 1(b) scatter plot of Bayesian correlation and Pearson correlation for all pairs of genes is shown. The Bayesian correlation was systematically lower. In Fig. 1(c) the distributions of gene expression of the top 3000 correlated pairs for Bayesian and Pearson correlations are shown. This result shows that the Bayesian correlation algorithm tends to identify correlations in genes that are highly expressed, compared with the correlations identified by Pearson correlation that identifies correlations in low expressed genes. We observed some low expressed genes among the Bayesian correlations, showing that Bayesian correlation is not equivalent to a higher threshold.

**Figure 1:**
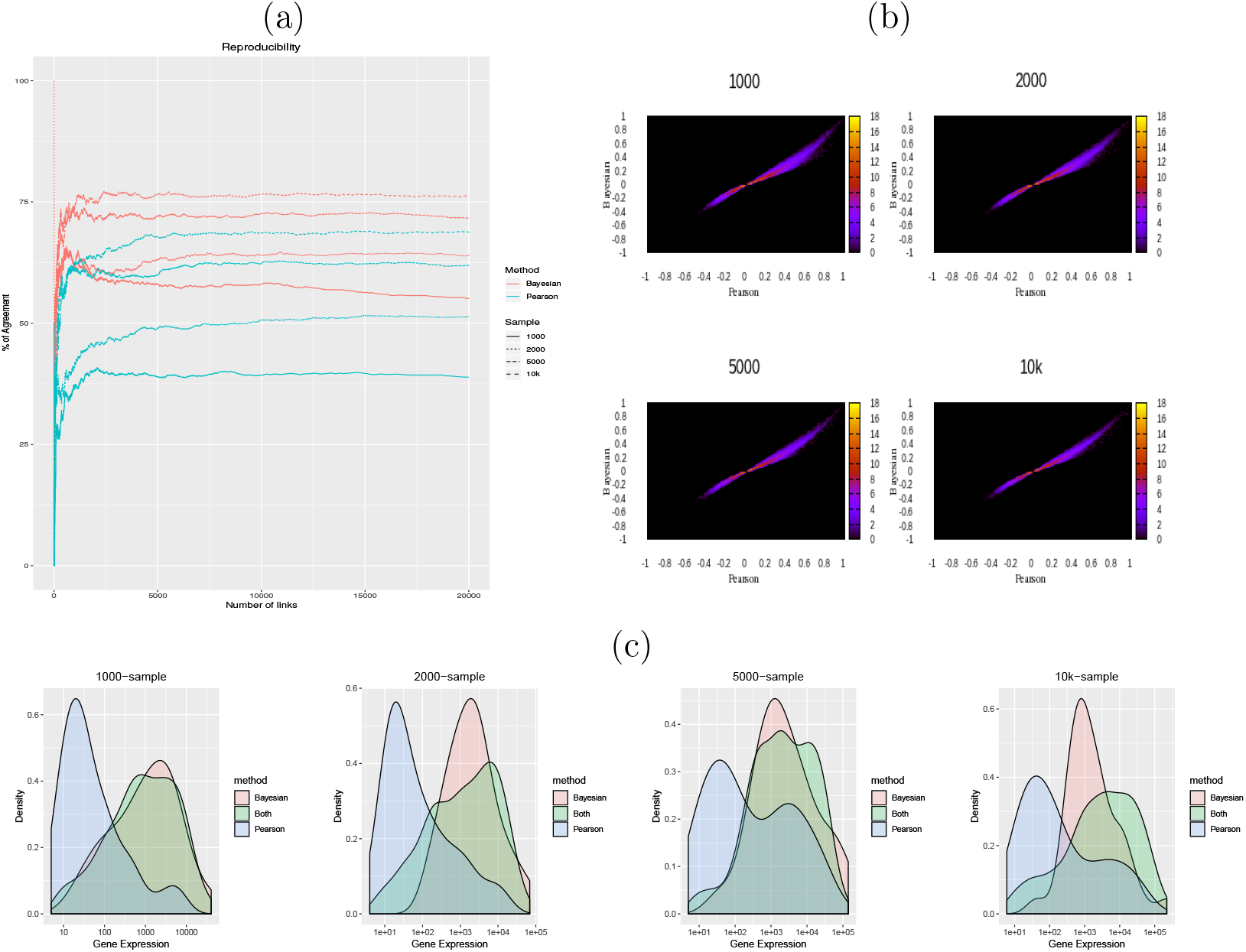
All cells correlation. (a) Percentage of pairs of correlated genes found in the two random samples as a function of the number of links sorted by correlation include repetitions (A correlated with B and B correlated with A are both included). (b) Scatter plot of the the Pearson correlation (x-axis) and Bayesian correlation (y-axis) for all genes, colored by the logarithm of the density. (c) Histogram of the total expression of the genes found in the top 3000 links.

### 3.2 Bayesian correlation is more robust than Pearson correlation to study similarity within small populations

We have shown a pronounced discrepancy between Pearson and Bayesian when the number of input cells is small. This result motivated us to study correlations within the different populations found in our data (Appendix A).

In this section, we restricted our analysis to the hepatocyte fraction. There are 33, 58, 111 and 200 hepatocytes in the 1000-sample, 2000-sample, 5000-sample and 10k-sample respectively.

In the hepatocytes clusters, the number of total UMIs was small, because of the small number of cells. That would result in thousands of 1 correlation in Pearson. For this reason, we used the dropout corrected data.

In Fig. 2(a) we observed that the reproducibility for correlations within small clusters is much higher with the Bayesian correlation algorithm than with Pearson. This difference was more pronounced for a small sample size, with around 90% of irreproducible results for Pearson in the 1000-sample and 2000-sample scenarios. The Bayesian correlation was systematically lower compared with Pearson (Fig. 2(b)). This effect decreased with the increase in the number of input cells. The distribution of the expression of the genes in the top 3000 correlated pairs for Bayesian and Pearson correlations is shown in Fig. 2(c). Pearson identified correlations in low expressed genes that are not considered by the Bayesian method.

**Figure 2:**
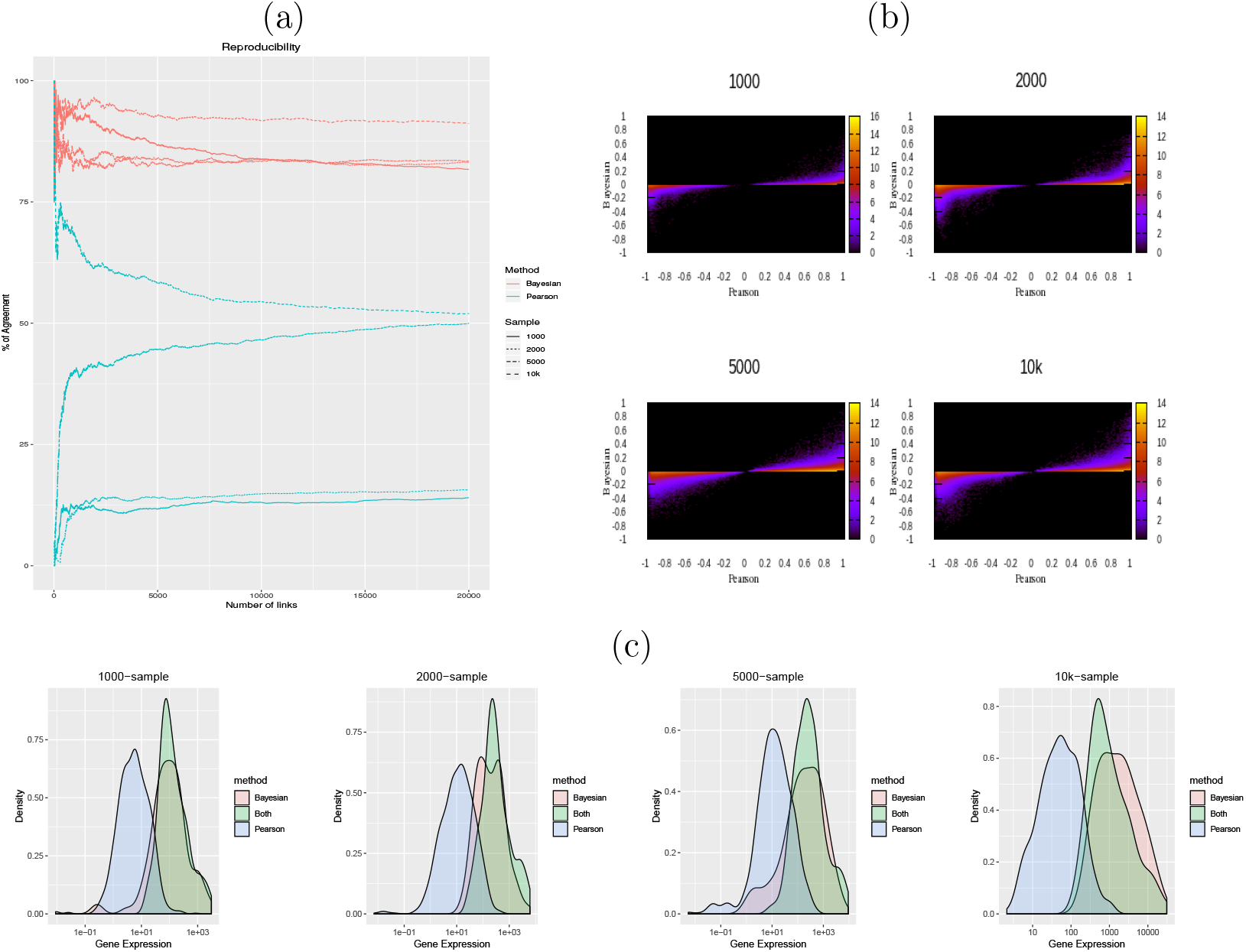
In cluster correlation: example of hepatocytes (a) Percentage of pairs of correlated genes found in the two random samples as a function of the number of links sorted by correlation. (b) Scatter plot of the the Pearson correlation (x-axis) and Bayesian correlation (y-axis) for all genes, colored by the logarithm of the density. (c) Histogram of the total expression of the genes found in the top 3000 links.

### 3.3 Bayesian correlation is more robust than Pearson correlation to study cluster similarity

Single cell sequencing allows the study of cells individually. However, combined with clustering techniques, it is possible to obtain bulk-like RNA-seq samples from pure populations. In this section, the single cells are grouped to study Bayesian Cluster correlation.

To test the reproducibility, the UMI matrix was split into two data sets and then the transformation of Eq. 2.9 was applied.

We showed that the reproducibility of our method is higher when the Bayesian method is applied. The agreement between Pearson correlations and Bayesian correlations increased with the number of cells (Fig 3. (a)). In Fig. 3(b) scatter plots of the Bayesian correlation (y-axis) vs the Pearson correlation (x-axis) are shown. The Bayesian correlation was systematically lowered. In Fig. 3(c) the distributions of the gene expression of the genes in the top 3000 correlated pairs for Bayesian and Pearson correlations are shown. As in the previous sections, the genes identified only by Pearson correlations were low expressed.

**Figure 3:**
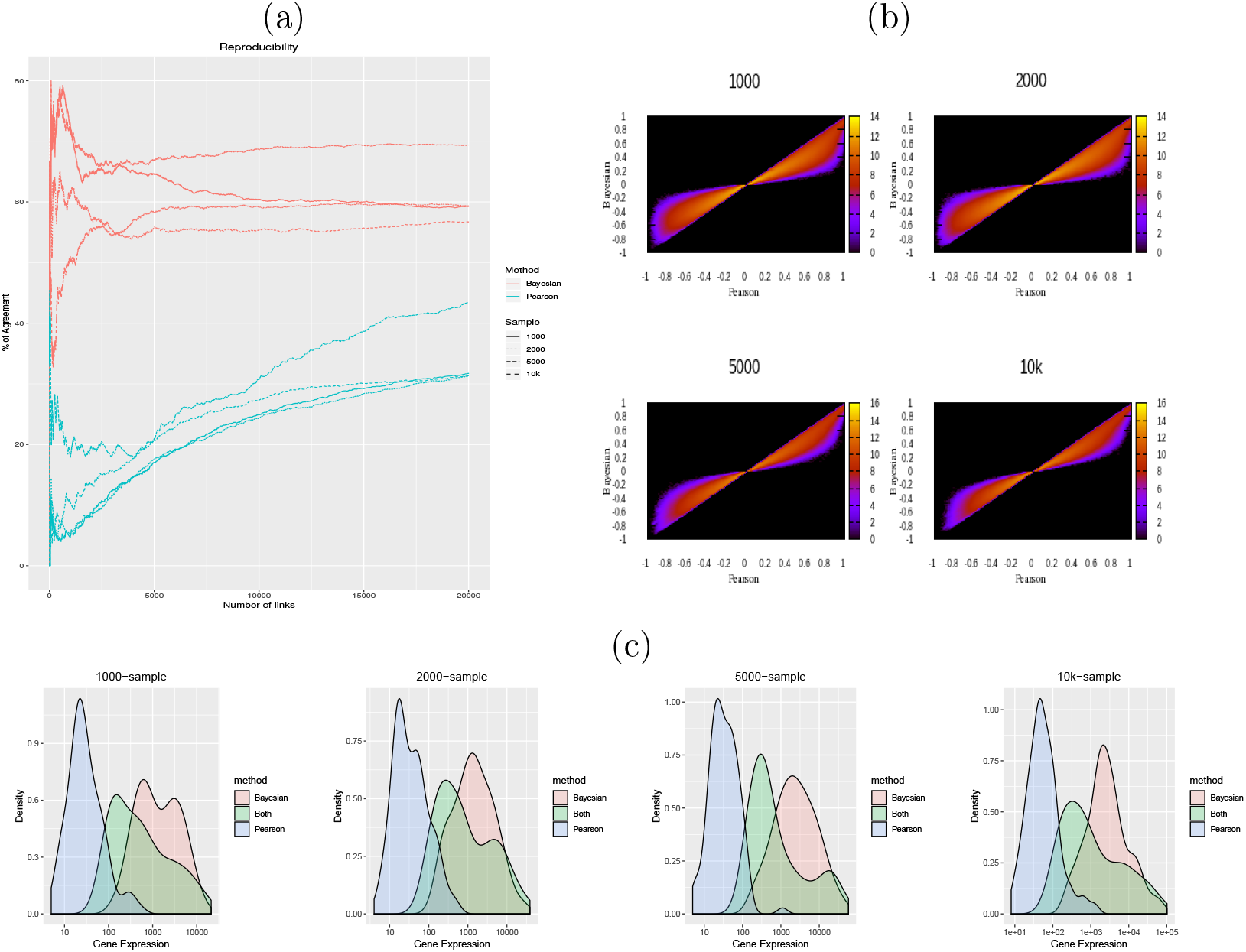
Cluster correlation. (a) Percentage of pairs of correlated genes found in the two random samples as a function of the number of links sorted by correlation. (b) Scatter plot of the the Pearson correlation (x-axis) and Bayesian correlation (y-axis) for all genes, colored by the logarithm of the density. (c) Histogram of the total expression of the genes found in the top 3000 links.

### 3.4 Robustness of Bayesian method for a varying number of cells

Thus far, to study the reproducibility of our method, we have compared it with the Pearson correlation by splitting each of our datasets in two groups. To determine the importance of the number of cells in an experiment, we next studied the agreement between our different samples.

In the previous sections, a strong disagreement between Bayesian and Pearson was observed in the Cluster Correlations and in In Cluster Correlations. For this reason, we restricted our analysis to those two scenarios.

We first restricted our analysis the cluster identified as the hepatocyte population with the MAGIC corrected data. Fig 4(a) shows an agreement between the different samples of around 50% for the top 20000 links with the Bayesian method. On the contrary, Pearson correlation showed a low agreement between samples, which was close to 0%.

**Figure 4:**
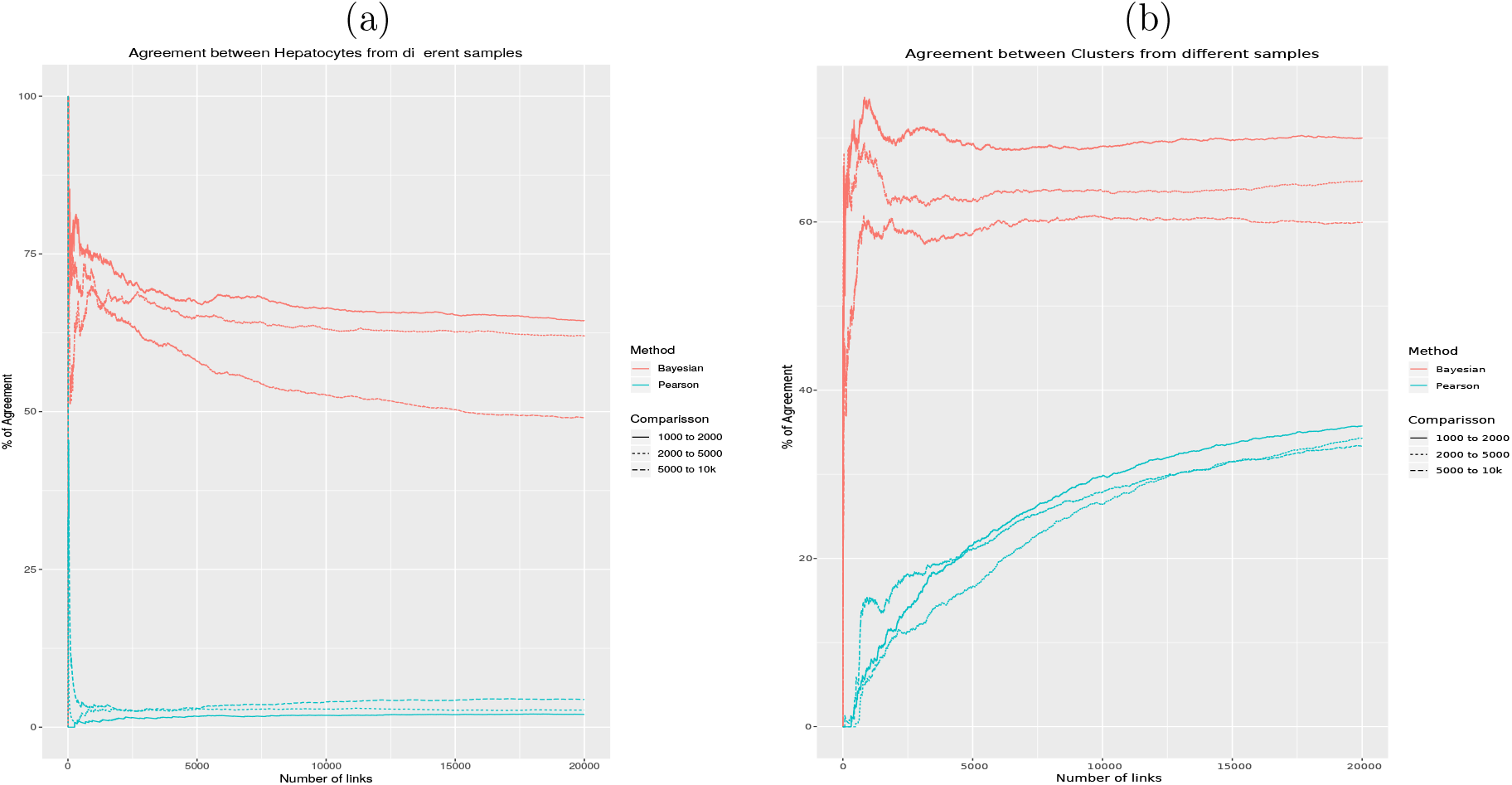
Percentage of correlated pairs found in the different samples using the Bayesian (red line) and the pearson (blue line) as a function of the number of links considered sorted by correlation. (a) Correlation in Hepatocytes. (b) Correlation in all clusters.

Second, we transformed our samples in bulk-like samples by means of Eq.(7). In Fig. 4(b) we observed that the agreement between samples is around 60% with the Bayesian method when 1000 links are considered. For the Pearson correlation, the agreement between the samples is smaller and it reaches 30% when 10000 links are considered.

### 3.5 Bayesian in cluster correlation assign high values to cell population markers

We have shown that Bayesian correlation increases the reproducibility by assigning low correlations to low expressed genes. In this section, we demonstrate that the genes present among the most highly correlated pairs of genes are biological meaningful.

In order to investigate the biological meaning of the correlated pairs found with our Bayesian method a set of hepatocyte markers from PanglaoDB [19] was downloaded on April 17th 2019. Fig. 5 shows the percentage of genes in the top correlations that are in this public database as a function of the number links considered for the different clusters with in cluster correlation. For the 4 samples, one cluster contained more genes of the database among the top correlated links than the others. In the 4 cases, that cluster was the one identified as the hepatocyte cluster.

**Figure 5:**
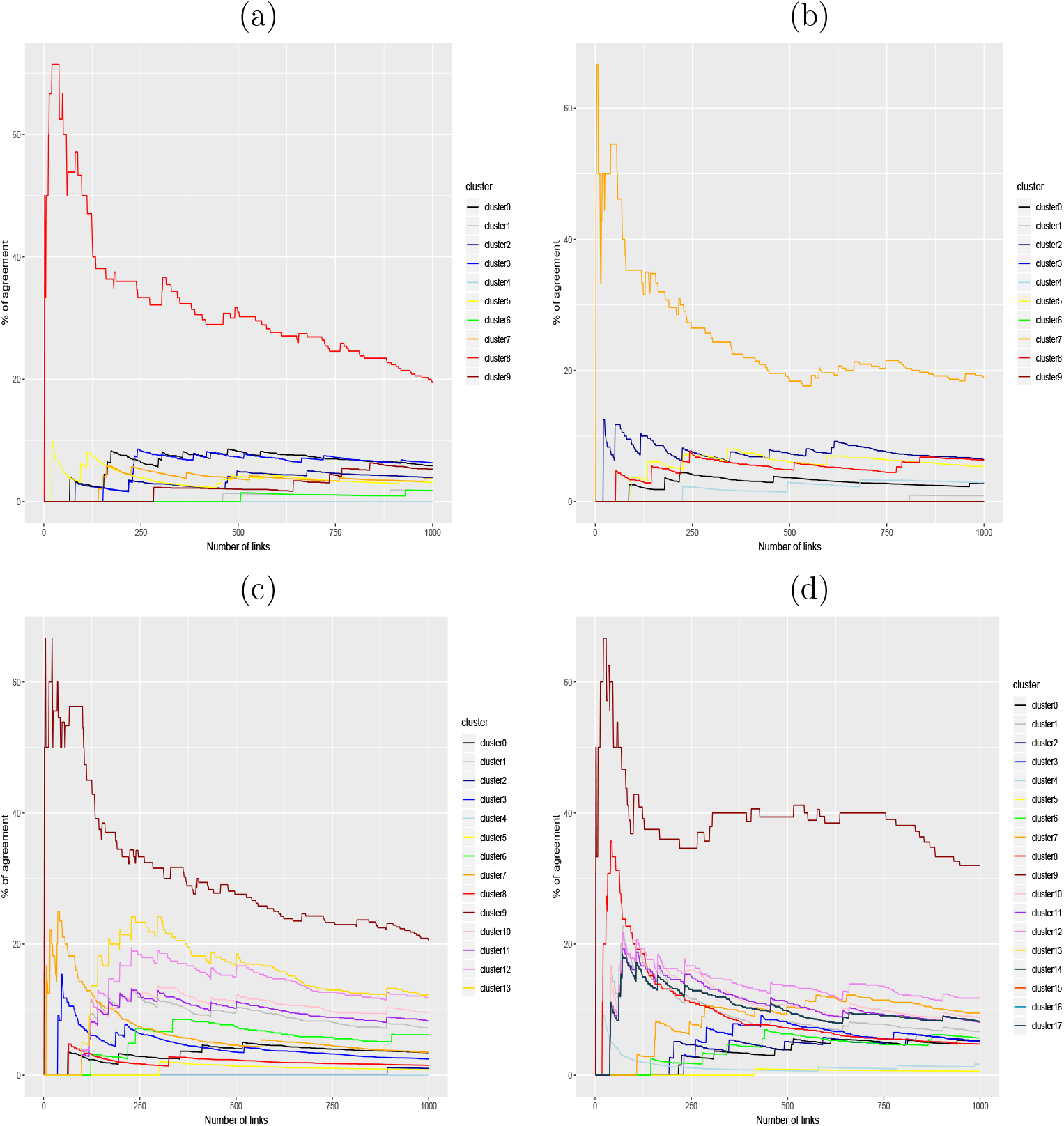
(a) 1000-sample, (b) 2000-sample, (c) 5000-sample, (d) 10k-sample. Percentage of genes in the top correlated pairs that are in the hepatocytes marker list from PanglaoDB as a function of the number of links considered sorted by Bayesian correlation. The correlation is computed as in a cluster correlation for each cluster independently identified with Seurat.

We have shown that the Bayesian correlation can be used to identify cell populations by looking at the genes present in the top correlated pairs. To investigate further the performance of this identification method, we compared it with two analogous methods. The first method is the same method using Pearson correlation. The second method is to intersect the markers obtained with Seurat with the hepatocyte marker list.

In order to make a fair comparison, when N genes are considered with the latter identification method, we choose the number of links that results in N unique genes. Fig. 6(a) showed that for a small number of genes the Bayesian correlation algorithm selected more hepatocytes markers than Seurat or Pearson correlation. For a larger number of cells Fig. 6(b,c,d) the Bayesian correlation and Seurat showed a similar performance and both are higher than Pearson correlation. When a large number of genes (e.g. 100 genes) is considered the three methods show a similar performance.

**Figure 6:**
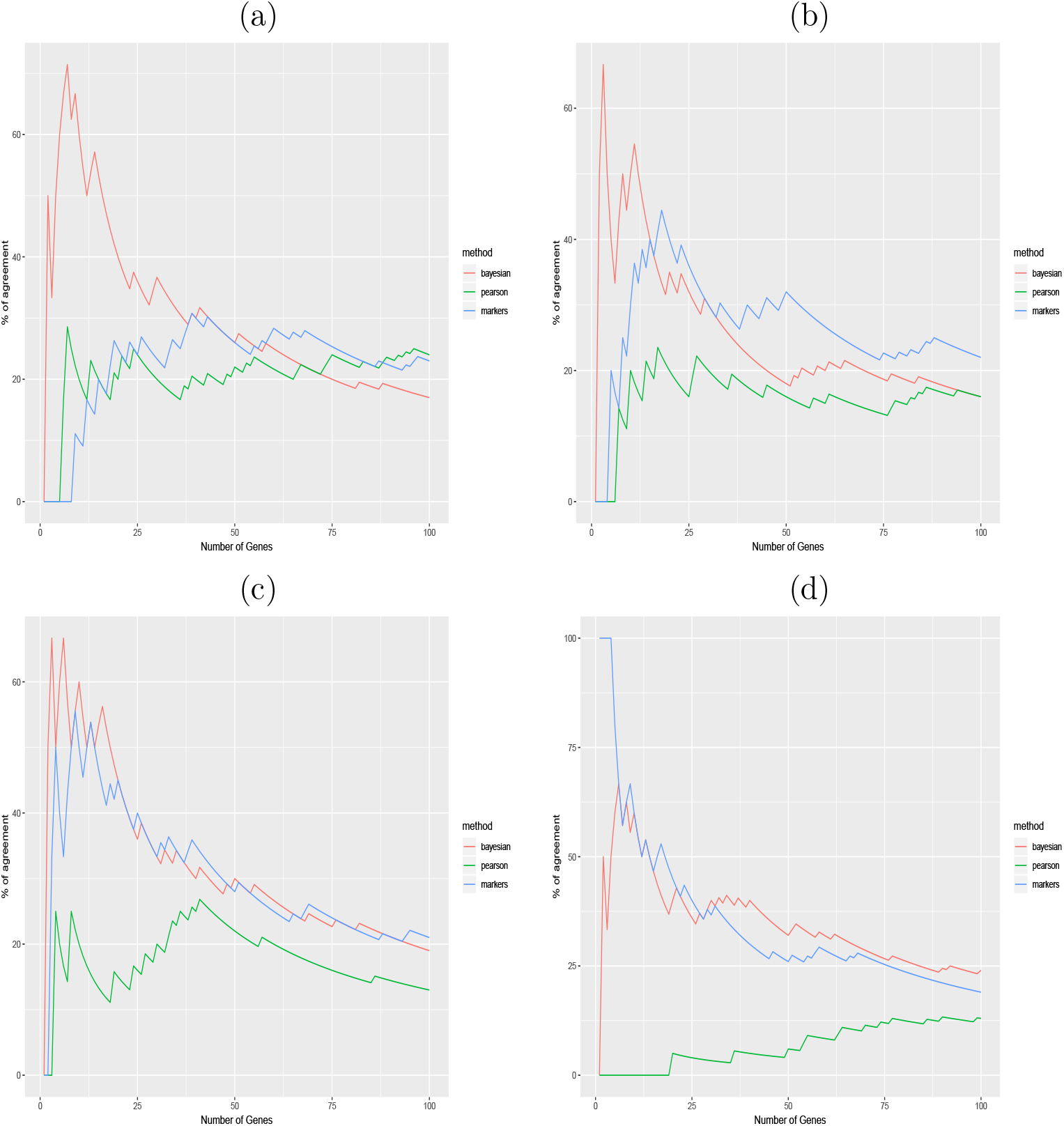
(a) 1000-sample, (b) 2000-sample, (c) 5000-sample, (d) 10k-sample. Percentage of genes in the top correlated pair in the Hepatocytes marker list from PanglaoDB as a function of the number of genes considered in the top pairs sorted by correlation (red and green) and by p-value in the markers identified by Seurat (blue).

## 4 DISCUSSION

We have presented a similarity measure of genes in scRNA-seq data, which suppresses correlations from low expressed genes, by extending the notion of Bayesian similarity from RNA-seq to scRNA-seq data. Our new Bayesian method allows scientists to study similarity between pairs of genes without discarding low expressed entities and avoiding biases. Thus, this new methodology is more resilient to noise and gives more reproducible results compared to Pearson. Moreover, the Bayesian scheme assigns high correlation to biologically relevant genes.

After splitting our samples in two groups we observed that the Bayesian correlation is more reproducible than Pearson correlation because it is not biased by low expressed genes. There was a more pronounced effect when the number of input cells was small. This result suggests that the Bayesian method can be useful to study very heterogeneous and rare populations.

We studied the correlation within different populations found in our data and focused on the hepatocyte population. Restricting the same methodology to these cells, after correcting the dropout, we observed that the reproducibility of the Bayesian correlations was higher than Pearson correlation. As before, we have seen that this difference in the methods decreases as the number of input cells increases.

Since the All cells correlations is biased by the number of cells in a cluster, we decided to study the Cluster correlation by summing the gene expression of all cells for each gene. Clusters with low cell numbers have low total reads and therefore are less resilient to noise. Note that the Bayesian correlation accounts for the total number of reads when computing the correlation. Applying the same methodology to the clusters, we observed that Bayesian correlation are more reproducible and are not biased by low expressed genes compared to Pearson.

After studying the three notions of correlation in our different samples, we have compared the results of the different samples, by comparing 1000 to 2000, 2000 to 5000 and 5000 to 10k. We studied the correlation in hepatocytes and we found that the agreement between samples was around 50% for the Bayesian method and close to 0 for the Pearson correlations. Then, we studied the cluster correlation and we showed that the samples agree more with the Bayesian correlation than with Pearson correlation.

Low expressed genes were detected in the top correlated genes using Bayesian correlations, therefore the method is not equivalent to a threshold.

To study the biological relevance of the correlated pairs identified by Bayesian correlation, we compared them with genes specific for hepatocytes from PanglaoDB. We have shown that the genes among the top correlated pairs tend to include more markers than Pearson correlation. Interestingly, when the number of cells is small, the performance of the method to identify markers is higher than the performance of Seurat [10]. There are two things that explain this fact. First, Bayesian correlation assigns higher values to high expressed genes, since markers are highly expressed they appear more. Second, genes with a specific functionality tend to be correlated with other genes important for that function since the pathways that activate them act on the entire population. Moreover, there are different types of hepatocytes (e.g. periportal and pericentral) and in those the expression of the markers is correlated. The optimal way to modify the Bayesian correlation algorithm for cell population identification, as well as a performance comparison with other identification methods such as SingleR [5] is left for future work.

We will further extend the Bayesian notion of similarity to mass cytometry [6] and CITE-seq [67]. By combining transcriptomic and proteomic analytical tools, we will build clustering methods to merge and validate results from single cell omic datasets. Development of a pipeline that integrates transcriptomic and proteomic data will clearly allow synergistic effects that cannot be identified by studying data independently.

Although our work mainly addresses bioinformatics questions, the dataset we have generated can be very useful for experiment design for liver researchers. Firstly, we provide a dataset of healthy mouse liver sequenced with high coverage. Second, our results show which cell populations can be identified within a scRNA-seq experiment and how many input cells need to be used.

Taken together, our results show that results from Bayesian correlations are more reproducible than results from Pearson correlations and have a higher biological relevance for analysis of scRNA-seq. Moreover, the number of sequenced cells have a small influence in Bayesian correlation results compared with Pearson correlation. Therefore, Bayesian correlation is a more robust measure of similarity for pairs of genes in scRNA-seq.

## DATA AVAILABILITY

All data is available on genome expression omnibus reprository with the GEO accession number GSE134134.

## ACKNOWLEDGEMENTS

We would like to thank the Swiss National Science foundation for funding (grant number 166594 and 173157) and the Unibe Initiator Grant (grant number 39027). We would like to thank the Next Generation Sequencing Platform of the University of Bern for performing the high-throughput sequencing experiments.

## A Cell type identification

During the preparation of this bioinformatics manuscript we created a dataset that can be useful for liver researchers. Our main goal is far from unravelling liver dynamics, however, we consider helpful for liver researchers to have an analysis of the biological samples. In this section, we describe the populations identified by Seurat clustering as well as the markers we use to classify them.

### A.1 scRNA-seq samples allow the identification of multiple parenchymal and non-parenchymal cell populations

We have identified 10,10,14 and 18 different cell populations in our 1000-sample, 2000-sample, 5000-sample and 10k-sample respectively. The multiple populations are showed in Fig. 7

**Figure 7:**
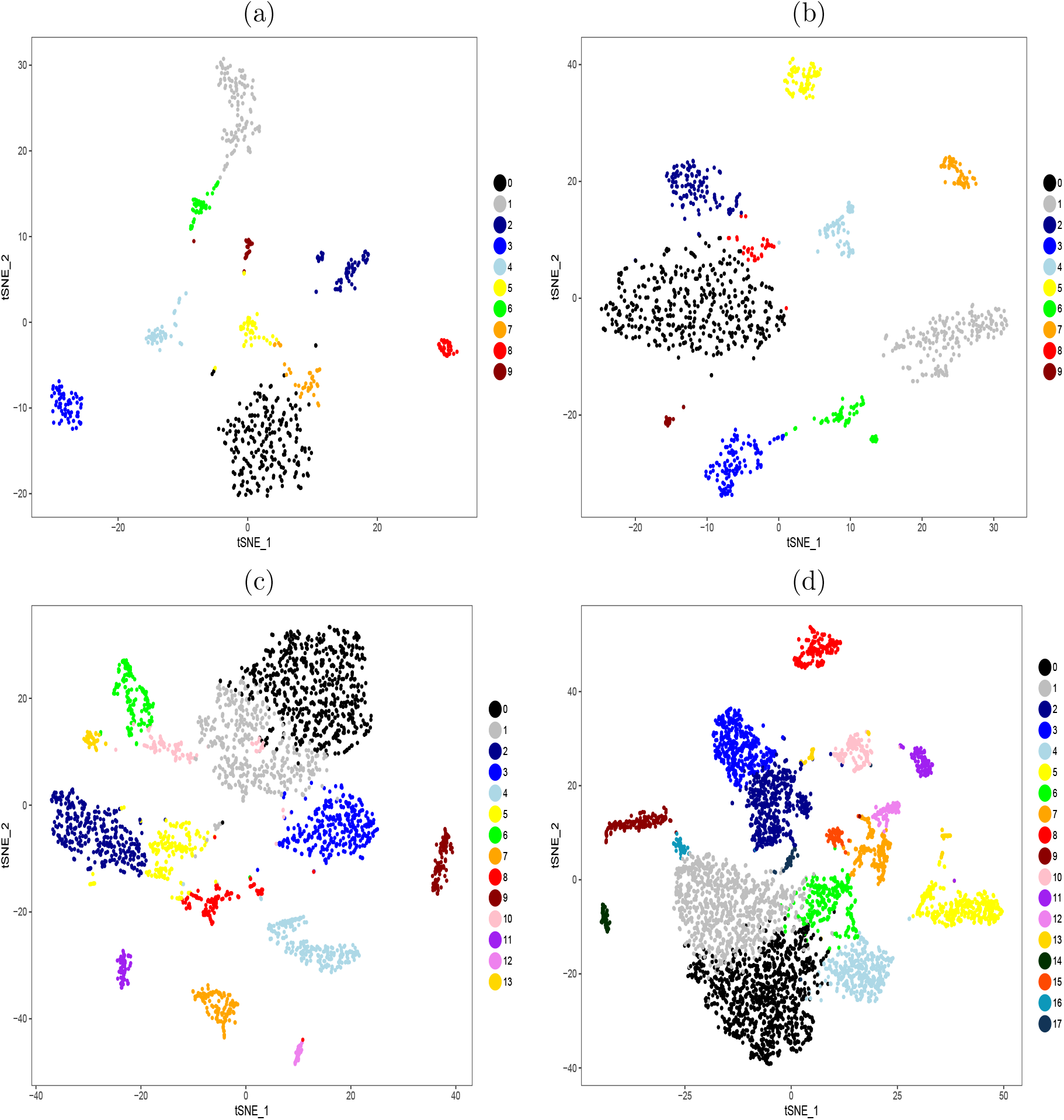
t-SNE visualization of our data, cells are colored by the cluster they belong to. (a)1000-sample (b) 2000-sample (c) 5000-sample. (d) 10k-sample.

### A.2 Cell types identified and markers

**1000-sample**

Unsupervised clustering of the 1000 cell sample identified 10 transcriptionally distinct populations. The different cell populations were identified using previously published markers form the literature.

- Cluster 0: Endothelial cells Markers: Pecam1,Ushbp1,Oit3,F8, Bmp2, C1qtnf1, Mmrn2, Pcdh12, Dpp4, Tek, S100a1, Tie1, Egfl7, Scarf1, Stab2, Lyve1, Icam2, Sox18, Flt4, Nr2f2 [59, 22, 26, 53, 57, 62, 28, 43, 54, 30, 18, 17, 15].
- Cluster 1: Macrophages (Kupffer cells) Markers: Adgre1, Csf1r, Cd163, Cd68, Marco, Vsig4, Irf7, Clec4f [35, 8, 46, 33, 38, 22].
- Cluster 2: Stellate cells Markers: Reln, Sparc, Col1a2, Rbp1, Des, Bmp5, Lrat [27, 2, 48, 38, 49, 28, 2, 16, 47].
- Cluster 3: Cholangiocytes Markers: Krt7, Krt19, Epcam, Sox9, St14 [38, 32].
- Cluster 4: NK/T-cells T-cell markers: Cd3g, Cd247, Gata3, CD28, Lat, Cst7, Cd3e, Cd4. NK cells markers: Nkg7, XcL1, CCl5, Cd7, Gzmb [38, 1, 44].
- Cluster 5: Endothelial cells Markers: Egfl7. Bmp2 Clec4g [22, 62, 79].
- Cluster 6: Dendritic cells Markers: Itgax Xcr1, Flt3, Cd24a, Ccr2, Clec9a [65, 50, 1, 8, 76].
- Cluster 7: Endothelial cells Markers: Pecam1, Ushbp1, Mmrn2, Tek, Flt4, Nr2f2 [22, 26, 53, 57, 28, 17, 15].
- Cluster 8: Hepatocytes Markers: Alb, Ass1, Cyp2f2, Asgr1, Apoa1, Mup3, Pck1, G6pc [22, 38].
- Cluster 9: Immune cells of Lymphoid branch Markers: Siglech, Ly6d, Runx2 [58, 61, 77, 42].

**2000-sample**

Unsupervised clustering of the 2000 cell sample identified 10 transcriptionally distinct populations. The different cell populations were identified using previously published markers form the literature.

- Cluster 0: Endothelial cells Markers: Pecam1, Ushbp1, Oit3, F8, Bmp2, Mmrn2, Pcdh12, Dpp4, Tek, S100a1, Scarf1, Stab2, Lyve1, Icam2, Sox18, Egfl7, Flt4, Nr2f2, Tie1 [59, 22, 26, 53, 57, 62, 28, 43, 54, 30, 18, 17, 15].
- Cluster 1: Macrophages (Kupffer cells) Markers: Adgre1, Csf1r, Cd163, Vsig4, Marco, Cd68, Cd5l, Irf7, Clec4f [35, 8, 46, 38, 33, 22].
- Cluster 2: Endothelial cells Markers: Clec4g, Egfl7, Bmp2, Oit3, Mmrn2 [22, 62, 79].
- Cluster 3: NK/T-cells T-cells markers: Cd3g, Cd247, CD28, Lat, Cst7, Cd3e. NK-cells markers: Nkg7, Xcl1, CCl5, Cd7, Gzmb [38, 1, 44].
- Cluster 4: Stellate cells Markers: Hhip, Reln, Sparc, Col1a2, Rbp1, Des, Lrat [27, 2, 48, 38, 49, 28, 16, 40].
- Cluster 5: Cholangiocytes Markers: Krt7, Krt19, Sox9, Epcam, Muc1 [38, 32, 45].
- Cluster 6: Dendritic cells Markers: Xcr1, Flt3, Cd24a, Ccr2, Clec9a, Itgax [50, 1, 8, 76, 65].
- Cluster 7: Hepatocytes Markers: Alb, Hnf4a, Ass1, Cyp2f2, Cyp2e1, Asgr1, Apoa1, Mup3, Pck1, G6pc [22, 38].
- Cluster 8: Endothelial cells Markers: Pcdh12, Sox18, Nr2f2 [22, 18, 15].
- Cluster 9: Immune cells of the lymphoid branch Markers: Ly6d, Sell, Cd19, Ms4a1, Ltb, Cd37 [4, 38, 1, 42].

**5000-sample**

Unsupervised clustering of the 5000 cell sample identified 14 transcriptionally distinct populations. The different cell populations were identified using previously published markers form the literature.

- Cluster 0: Endothelial cells Markers: Pecam1, Ushbp1, Oit3, F8, Bmp2, Pcdh12, Dpp4, Tek, S100a1, Scarf1, Stab2, Lyve1, Icam2, Sox18, Egfl7, Flt4, Nr2f2, Tie1 [59, 22, 26, 53, 57, 62, 28, 43, 54, 30, 18, 17, 15].
- Cluster 1: Endothelial cells Markers: Pecam1, Ushbp1, Oit3, F8, Bmp2, Mmrn2, Pcdh12, Dpp4, Tek, S100a1, Stab2, Lyve1, Icam2, Sox18, Egfl7, Flt4, Nr2f2, Tie1 [59, 22, 26, 53, 57, 62, 28, 43, 54, 30, 18, 17, 15].
- Cluster 2: Macrophages/Kupfercells Markers: Adgre1 Csf1r, Cd163, Vsig4, Marco, Cd68, Cd5l, Irf7, Clec4f [35, 8, 46, 33, 38, 22].
- Cluster 3 Endothelial cells: Markers: Pecam1, Oit3, F8, Bmp2, Mmrn2, S100a1, Icam2, Egfl7, Clec4g [59, 22, 26, 53, 62, 28, 54, 30, 18, 17, 15, 79].
- Cluster 4: T and NK cells T-cell markers: Cd3g, Cd247, Trac, CD28, Lat, Cst7, Cd3e. NK-cell markers: Nkg7, Xcl1, CCl5, Cd7, Gzmb [38, 1, 44].
- Cluster 5 Macrophages: Markers: Clec4f Csf1r, Cd163, Vsig4, Marco, Cd68, Cd5l [8, 46, 33, 38, 22].
- Cluster 6: Stellate cells Markers: Hhip, Reln, Sparc, Col1a2, Rbp1, Des, Lrat [27, 2, 48, 38, 49, 28, 40, 47].
- Cluster 7: Cholangiocytes Markers: Krt7, Krt19, Sox9, Epcam, Muc1, St14 [38, 32, 45].
- Cluster 8: Dendritic cells Markers: Xcr1, Flt3, Cd24a, Ccr2, Clec9a [50, 1, 8, 76].
- Cluster 9: Hepatocytes Markers: Alb, Ass1, Cyp2f2, Cyp2e1, Asgr1, Apoa1, Mup3, Pck1, G6pc [22, 38].
- Cluster 10: Endothel cells: Markers: Pecam1, Oit3, F8, Bmp2, Mmrn2, Dpp4, Tek, S100a1, Stab2, Sox18, Egfl7, Flt4, Nr2f2, Tie1 [59, 22, 26, 53, 57, 62, 28, 54, 18, 17, 15].
- Cluster 11: B-cells Markers: Cd19, Ms4a1, Ltb, Cd37, Cd22, Cd79a, cd79b, Cd69 [1, 4, 38, 72].
- Cluster 12 Immune cells of the lymphoid branch Markers: Siglech, Ly6d, Runx2, Klra17 [58, 61, 77, 42].
- Cluster 13: Unknown

**10k-sample**

Unsupervised clustering of the 10k cell sample identified 17 transcriptionally distinct populations. The different cell populations were identified using previously published markers form the literature.

- Cluster 0: Endothelial cells Markers: Clec4g, Pecam1, Tek [59, 26, 79].
- Cluster 1: Endothelial cells Markers: Clec4g, Pecam1, Dpp4, Lyve1 [22, 43, 59, 79].
- Cluster 2: Macrophages Markers: Csf1r, Adgre1, Cd163, Vsig4, Marco, Cd68, Cd5l, Irf7, Clec4f [35, 8, 46, 46, 33, 38].
- Cluster 3: Macrophages Markers: Adgre1, Csf1r Cd163 Vsig4 Marco Cd68 Cd5l Irf7 Clec4f [8, 33, 35, 38, 46].
- Cluster 4: Endothelial cells Markers: Oit3, Bmp2, Mmrn2, Icam2, Sox18, Flt4, Clec4g, Egfl7 [22, 53, 57, 62, 28, 54, 30, 18, 17, 15, 79].
- Cluster 5: NK-cells/T-cells T-cell markers: Trac, Cd3g, Cd2. NK-cells markers: Nkg7, Xcl1, CCl5 [38, 1, 44].
- Cluster 6: Endothelial cells Markers: Pecam1, Ushbp1, Bmp2, Stab2, Egfl7, Clec4g [22, 53, 57, 62, 28, 54, 30, 18, 17, 15, 79, 59].
- Cluster 7: Stellate cells Markers: Hhip, Reln, Sparc, Rbp1, Des, BMp5, Pdgfrb, Lrat, Hand2 [27, 2, 48, 38, 49, 28, 16, 9, 41, 25, 40, 47].
- Cluster 8: Cholangiocytes Markers: Krt7, Krt19, Sox9, Epcam, Muc1, St14 [38, 32, 45].
- Cluster 9: Hepatocytes Markers: Alb, Apoa1, Mup3, Ass1, Cyp2f2, Cyp2e1, Asgr1, Pck1, G6pc [22, 38].
- Cluster 10: Dendritic cells Markers: Xcr1, Ccr2, Itgax, Flt3, Cd24a, Ccr2 [65, 50, 1, 8, 76].
- Cluster 11: Immune cells: B cells Markers: Cd19, Ms4a1, Ltb [1, 4, 38].
- Cluster 12: Stellate cells Markers: Hand2, Hhip, Sparc1, Des, Reln, Rbp1 [27, 2, 48, 38, 49, 28, 40].
- Cluster 13: Unknown
- Cluster 14: Immune cell from the lymphoid branch Markers: Siglech, Runx2, Klra17 [58, 61, 77, 42].
- Cluster 15: Stellate cells Markers: Pdgfrb, Lrat, Hand2, Hhip, Reln, Sparc, Des, Rbp1 [27, 9, 2, 48, 38, 49, 28, 40].
- Cluster 16: Unknown
- Cluster 17: Endothelial cells Markers: Ptprb, Pecam1 [66, 59].

